# Out of the cradle: expansion from the natal ecosystem erodes morphological specificity in endemic Kamchatkan charrs (genus *Salvelinus*)

**DOI:** 10.64898/2026.09.03.749089

**Authors:** Evgeny Esin, Ludmila Zinevich, Kirill Kuzishchin, Marina Gruzdeva, Grigorii Markevich

**Author notes:** **Corresponding author:** Esin, E.V.

## Abstract

The contribution of postglacial adaptive radiations to maintaining regional biodiversity of freshwater fish remains debatable. To uncover whether such radiations can lead to the emergence of new populations in neighbouring ecosystems, we have reconstructed the evolutionary history of the endemic *Salvelinus malm*a assemblage from Lake Kronotskoe in Kamchatka. We traced the dispersal of endemic evolutionary lineages from the lake into neighbouring basins and quantified the morphological divergence between the lacustrine populations and their derivatives. To this end, we first examined the phylogeny, genetic differentiation, current hybridization and historical gene flow of eleven charr groups using genome-wide SNPs, followed by a charr morphometric comparison. We found that the Lake Kronotskoe lineages of the predatory and littoral benthivorous charrs invaded small peripheral waterbodies via temporary riverbed interconnections during the Holocene and continue to migrate downstream from the landlocked ecosystem, hybridizing with the anadromous *S. malma*. These dispersal processes have resulted in three new isolated populations and one hybrid anadromous population. The evolution of morphotypes in derived populations occurred independently of their phylogenetic history, resulting in the generalized morphology that is characteristic of pelagic or riverine dwellers. These findings show that the postglacial expansion of adaptive morphs from the cradle ecosystem bolsters regional diversity, and that migrants quickly lose ancestral adaptive traits, forming new morphotypes in response to changing environmental pressures.

## Introduction

Adaptive radiation is a rapid evolutionary diversification of a single phylogenetic lineage into many new adaptive forms that share the ecological opportunities within an ecosystem (Bird et al., 2012; Marques et al., 2019; McGee et al., 2020; Stroud et al., 2025). From a biodiversity perspective, adaptive radiations account for a significant proportion of species lists (or biodiversity units of another status) (Stroud et al., 2025). This phenomenon is clearly manifested in freshwater fish, with iconic examples include the cichlid fauna of African Great lakes (Seehausen, 2015) and silversides from the ancient lakes of Sulawesi (von Rintelen et al., 2012). The complexity of sympatric assemblages resulting from fish radiations depends on the ancestor genome’s heterogeneity, the ecosystem characteristics, and the age of the radiation process (Recknagel et al., 2017; Doenz et al., 2019; Blain et al., 2023; Tiddy et al., 2024). The longer an assemblage exists, the more complex its evolutionary processes become and the deeper the specialization in specific niches may develop (Sexton et al., 2017). The ultimate outcome of the adaptive radiation is an endemic assemblage of specialized species/ecomorphs confined to a particular waterbody. This tight coupling of diversification to a specific environment raises the question: do locally radiated endemics play a role in maintaining global biodiversity, and are they so specialized to maintain specific adaptive traits outside their native ecosystem?

Certain endemic species, originating from ancient radiations, are documented to have gained the opportunity to expand from their natal ecosystems to the neighbouring water systems. These events may have occurred during periods of drastic geological rearrangement or climate fluctuation. For example, endemic Cottidae of Lake Baikal have occupied the outflowing Angara River and remote Yenisei tributaries (Sideleva et al., 2025). The descendants of Lake Tanganyika Cichlidae spread across remote basin tributaries during periods of high rainfall (Kullander et al., 2011). Specialized African scraper, benthivorous and predatory Cyprinidae settled in a number of watercourses in the Abyssinian highlands (Levin et al., 2019; 2020). A specific example is the troglomorphic Mexican tetras of the genus *Astyanax*, which are believed to have spread through flooded karst channels between caves (Borowsky, 2018). Collectively, these cases demonstrate that even highly specialized endemics can, under certain conditions, serve as sources for broader regional diversification. As evidenced by extant publications, the settlers retained at least some of the distinctive morphological features of their endemic ancestors.

Salmonids of the genera *Coregonus* and *Salvelinus* typically diversify into different adaptive morphs in postglacial lakes across the Holarctic (Skúlason et al., 2019). These ecomorphs are genetically differentiated groups that have acquired phenotypic differences beneficial for resource partitioning and ecological specialization (Streelman & Danley, 2003; Schluter & Conte, 2009; Hernández-Hernández 2019). There are several scenarios commonly considered for the origin of salmonid ecomorphs: sympatric diversification of a single ancestor within an ecosystem; invasion of several stocks into a lake, with each stock becoming ecologically specialized; and resettlement of specialized ecomorphs as a result of ecosystem fragmentation (Hudson et al., 2007; Turgeon et al., 2017; Skúlason et al., 2019). However, we found no documented examples of a successful resettlement of salmonid ecomorphs, having formed within a particular lake-river system, into another basin. Ecomorphs with a specialized morphotype and feeding behavior appear to be strictly confined to their cradle ecosystems. Even if an anadromous morph that systematically leaves the ecosystem does exist, it remains part of the endemic assemblage due to its strict reproductive philopatry, as demonstrated by charrs of the genus *Salvelinus* in Scandinavia, Greenland and Kamchatka (Nordeng, 2009; Doenz et al., 2019; Busarova, 2022).

Consequently, it remains unknown whether salmonid ecomorphs of sympatric assemblages spread and, if so, whether they retain specific adaptive features in new habitats, at least for a relatively short evolutionary period of the Holocene.

To investigate the possibility of salmonid ecomorphs to expand their range and transform their morphotypes in new habitats, we studied the assemblage of charrs in the Kronotskoe-Uzon catchment in Kamchatka. An explosive multi-step diversification of Dolly Varden *Salvelinus malma*, which now inhabits all rivers of the region, has occurred following its invasion into the mountain Lake Kronotskoe (Markevich et al., 2018; Esin et al., 2020). Niche-associated morphological differences (Markevich et al., 2018) and traces of selection detected in the genome (Woronowicz et al., 2024) suggest an adaptive basis for the Lake Kronotskoe radiation. Different genetic markers indicate the complex demographic history of ecomorphs from Lake Kronotskoe and neighbouring water systems (Esin et al., 2020; 2025). Ecomorphs vary in their swimming ability and tendency to migrate (Markevich et al., 2021). However, the phylogenetic and morphometric relationships of ecomorphs from the central lake and surrounding smaller waterbodies remain to be elucidated.

The modern ecosystem of Lake Kronotskoe began to form around 22‒24 thousand years ago, after the retreat of the Pleistocene glacier and the last major volcanic event in that area (Golub, 2006; Ponomareva et al., 2025). Crucially, several local geological events restructured the basin after the Lake Kronotskoe charr had diversified into its main lineages. The rapids on the Kronotskaya River were formed (Shantser & Melekestsev, 1967), the small crater lakes Krokur and Dal’nee originated (Ponomareva et al., 2017), and the Taunshits volcano erupted, causing huge landslides that altered the upper reaches of rivers in the southern part of the area (Ponomareva et al., 2006). These dramatic rearrangements of the water network could enable the Lake Kronotskoe charrs to resettle in the neighbouring basins and new crater lakes via intercepting tributaries and migrating downstream.

In this study, we traced the routes of dispersal of different evolutionary lineages of charrs from Lake Kronotskoe to the neighbouring Uzon and Kamchatka River basins, and then assessed the morphological variation between the endemic Lake Kronotskoe ecomorphs and the derived populations. To accomplish this, we examined the SNP-based phylogeny, genetic differentiation, current hybridization and historical gene flow, as well as the landmark-based morphometric differences between the charrs. Eleven samples from Lake Kronotskoe and the surrounding basins were used in the analysis. We hypothesize that the strict sedentarism of endemic charrs in the lake is unlikely. The ecomorphs are thought to have spread from the Lake Kronotskoe tributaries to the surrounding waterbodies (i); novel derivative populations have formed morphotypes that differ from the adaptive morphotypes of Lake Kronotskoe endemics (ii); endemic charrs have also migrated downstream from the landlocked catchment through rapids and formed new hybrid population(s) with Dolly Varden in the outflowing rivers (iii). Moreover, the presence of genetic markers and specific morphological characteristics in Dolly Varden populations inhabiting remote rivers is a subject of interest.

## Material and Methods

### Kronotskoe-Uzon charr diversity

Lake Kronotskoe is a large (246 km^2^) and deep (up to 136 m) waterbody, with its main tributaries inflowing from the south-west and draining the northern flattened slopes of the Uzon and Taunshits edifices (**Fig. 1a**). Steep mountain ridges to the north and west divide the tributaries of Lake Kronotskoe from the Kamchatka River basin. The outflowing Kronotskaya River passes through huge rapids, which are impassable for fish migrating upstream.

**Figure 1.**
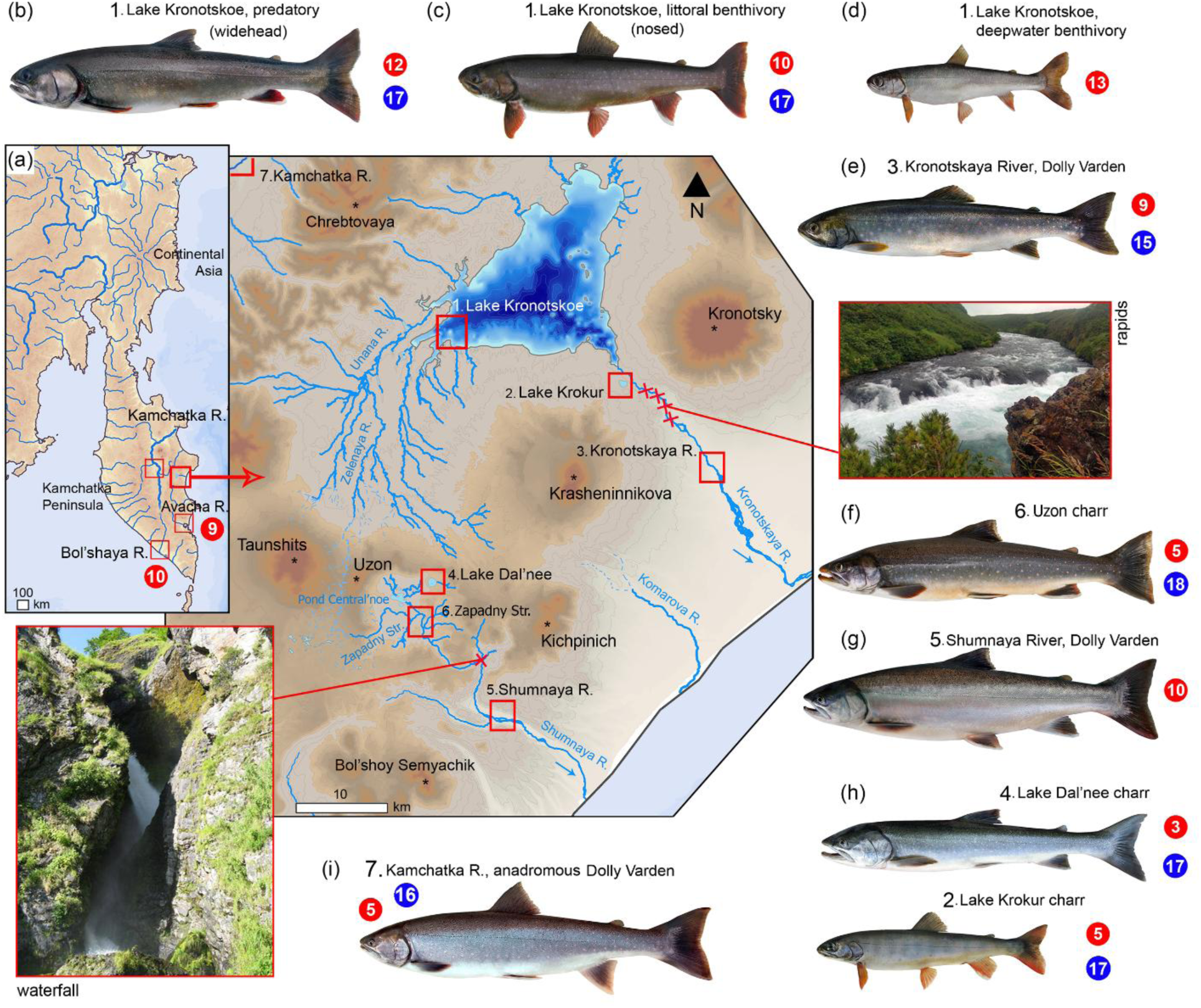
Scheme of the Kronotskoe-Uzon catchment area and its position on the Kamchatka Peninsula (**a**: the captioned squares indicate the sampling locations; symbols: * ‒ volcano peaks, x ‒ waterfalls and rapids, ···· ‒ temporary watercourses, → ‒ flow direction; photos show the obstacles in outflowing rivers). The charr groups used for the analysis (**b‒i**: around the scheme; the numbers in the adjacent red circles show the number of specimens used for the genome-wide analysis; blue circles ‒ specimens used for the morphometric analysis; the numbers in front of the sample name correspond to the sampling location on the catchment area scheme).

A recent genome-wide analysis revealed the presence of three evolutionary lineages within the Lake Kronotskoe charr assemblage (Woronowicz et al., 2024). According to trophic-ecological and morphological markers, these lineages represent large-sized piscivorous (predatory, **Fig 1b**), mid-sized littoral benthivorous (nosed, **Fig. 1c**) and small-sized deepwater benthivorous (**Fig. 1d**) charrs (Esin et al., 2024). The predatory charrs spawn in the upper reaches of the tributaries, the nosed charrs undertake shorter spawning migrations, and the deepwater charrs spend their entire lives in the hypolimnetic zone of the lake (Markevich et al., 2021). The Kronotskaya River downstream of the rapids is used for reproduction by the anadromous omnivorous Dolly Varden (**Fig. 1e**) (Esin et al., 2020).

The Uzon caldera, located to the south, is drained by the Shumnaya River, which drops over a waterfall 10 km from its source. The river is sourced by Pond Central’noe (3.0 km^2^ and 3 m deep) and the Zapadny Stream, which flows from the southern slope of the passage between the Uzon and Taunshits volcanoes (**Fig. 1a**). The Uzon waterbodies upstream of the waterfall are home to the endemic Uzon charr (**Fig. 1f**) (Esin et al., 2015), while the anadromous Dolly Varden reproduces in the lower reaches of the Shumnaya River (**Fig. 1g**).

The area’s drainage system has also been augmented by two small drainless lakes (both 1.3 km in diameter), which are filled maar craters, Dal’nee and Krokur (**Fig. 1**), formed in the mid-Holocene (Ponomareva et al., 2017). Both maars are inhabited by the isolated charr populations (**Fig. 1h**) (Esin et al., 2015; Esin & Markevich, 2019).

### Sample collection

In 2023, we sampled charrs using gill nets (mesh size 20‒32 mm) with permission № 041 dated 10.07.2023, issued by the Far Eastern Interregional Administration of Rosprirodnadzor. The most prevalent ecomorph of each endemic charr lineage was sampled in Lake Kronotskoe. These were the predatory widehead morph, the littoral nosed (N1) morph, and the deepwater smallmouth morph (Esin et al. 2020). These charrs differ in their specific phenotypes and have never been caught outside the Lake Kronotskoe basin. Landlocked charrs were also collected from the Zapadny Stream, Lake Dal’nee and Lake Krokur. Anadromous Dolly Varden was sampled in the Kronotskaya and Shumnaya rivers, which both drain the Kronotskoe-Uzon catchment. A total of 101 adult individuals from the Kronotskoe-Uzon drainage system, showing no pre-spawning changes of the head and body shape, were selected for morphological analysis. Of these, 67 individuals were subsampled for genetic analysis (**Fig. 1**).

Beyond the Kronotskoe-Uzon drainage system, five anadromous Dolly Varden were caught for genetic analysis in the Kamchatka River, beyond the mountain ridge from Lake Kronotskoe (**Fig. 1i**). The samples for genetic analysis were also sampled from 19 anadromous Dolly Varden in two remote rivers in the east and west of Kamchatka. According to Federal Law no. 166-FZ, fishing for the non-endemic *S. malma* does not require special permission.

### SLAF sequencing

The fin clips sampled for genetic analysis were preserved in 96% EP ethanol. High-molecular-weight DNA was extracted with the QIAamp DNA Mini Kit (Qiagen, Hilden, Germany) according to the manufacturer’s protocol. DNA concentration was measured using the Qubit 3.0 Fluorometer and the Qubit dsDNA HS Assay Kit (Invitrogen, USA), and DNA integrity was evaluated by electrophoresis on a 1% agarose gel.

For Specific-Locus Amplified Fragment sequencing (SLAF-seq) library preparation, an *in silico* pilot experiment was first conducted to select a combination of restriction enzymes and a fragment size range that would maximize genome coverage and uniformity while minimizing the repetitive sequences. Based on this design, genomic DNA was digested with *RsaI* and *HaeIII* (NEB, USA) to generate SLAF tags. The 3′ ends of the digested fragments were then adenylated (A-tailed) to facilitate adapter ligation.

Dual-indexed sequencing adapters were ligated to the A-tailed fragments. The adapter-ligated DNA was then PCR amplified, and the PCR products were purified using VAHTS DNA Clean Beads (N411-03, Vazyme, China). The purified PCR products were separated on a 2% agarose gel, and the DNA fragments within the target size range were excised and gel-purified. This size-selected DNA was subjected to a second round of PCR amplification, followed by another bead-based purification to produce the final SLAF-seq library. Completed libraries from 91 samples were quantified with a Qubit 3.0 fluorometer, and the fragment size distribution was verified using a Qsep-400 capillary fragment analyzer. Finally, the libraries were sequenced on an Illumina NovaSeq X Plus platform (Illumina, San Diego, CA, USA) to generate 150 bp paired-end reads, following the manufacturer’s instructions. Quality control tests were performed using FASTQC v0.11.8 (Andrews, 2010) and MultiQC v1.13 (Ewels et al., 2016).

### SNP calling and filtering

Paired, demultiplexed reads were filtered and adapter-trimmed using process_ragtags (v1.44) from STACKS (v2.68; Rochette et al., 2019) with the parameters: --filter-illumina -c -q -r --disable_rad_check. The filtered reads were mapped to the *Salvelinus* sp. genome ASM291031v2 (GeneBank accession number: GCF_002910315.2) using BWA MEM (v0.7.17-r118; Li & Durbin, 2009) with the mean success = 97.9% (the parameters: --M and -R to define read groups). The aligned reads were sorted, filtered and indexed with SAMtools (v1.6; Li et al., 2009). Variant calling was performed using the reference-based pipeline implemented in the ref_map.pl script from STACKS, with -r 0.50 --vcf --vcf-all for the initial run of ‘populations’. We flagged putative paralogs as loci showing H_obs > 0.6 and F_IS < -0.5 in all groups based on per-locus summary statistics from ‘populations’ (Stacks), and created a whitelist of the remaining loci. We then exported two filtered datasets using populations: a strict one (-r 0.70, one SNP per locus --write-single-snp, --vcf, --plink, --ordered-export, -W) for the admixture analysis, and a more permissive one (-r 0.50, all SNPs retained, --fstats, --vcf, --ordered-export, -W) for phylogenetic and introgression analyses.

In total, we obtained 1,590 million paired reads across 91 samples, with a mean of 11.0 million paired reads per sample. ‘Gstacks’ inferred genotypes using a maximum likelihood framework across 14,274,074 loci (mean length 229 bp), with 90.7% of diploid loci consistently phased. After PCR duplicate removal, the mean per sample coverage per locus was 5.4× with Std. Dev 1.6× (from 1.9× to 11.3×).

Although the low coverage, ’gstacks’ aggregated evidence across all samples at each locus for SNP discovery and remained reliable at low individual coverage (Rochette et al., 2019). Following population-level filtering (one sample removed; loci required to be genotyped in ≥50% of samples per population, -r 0.5), 5,444,965 loci were retained (mean genotyped length 206 bp), with 17,923,592 variant sites. For downstream phylogenetic analysis, we additionally filtered 1,673 loci with excess heterozygosity and strongly negative Fis (putative paralogs), leaving 17,772,739 variant sites, for population structure analyses, we retained one SNP per locus using the --write-single-snp option in ‘populations’, which yielded 14,768,687 SNPs.

### Phylogeny

The phylogenetic relations of the charrs were analyzed using the Maximum Likelihood (ML) approach in IQ-TREE 2 (v2.1.4; Minh et al., 2020), the model TVMe+R3 was applied (parameters: --m MFP for model choice, then --bb 1000 -alrt 1000 -st DNA). Multiple sequence alignment of SNPs was created using the -- phylip-var option of the ‘populations’, with retention of loci genotyped in at least 80% of all samples and SNPs with a minor allele count of three (--min-mac 3). To take into account the absence of constant sites in the alignment, an ascertainment bias correction (+ASC) model (Lewis, 2001) was applied to all substitution models in a best-fit model selection process performed with model finder. The bootstrap node support was based on 1,000 replicates. The tree was rooted to the ASM291031v2 genome of *S. malma* from the Tree River. As the reference genome was positioned inside one of the clades, the tree was re-rooted using ape (v5.3; Popescu et al., 2012) in R (v3.5.0; R Core Team). The resulting tree was visualized using ggplot2 (v3.4.2), ape(v5.3), phytools (v 0.7.80) and ggtree (v1.14.6) in R.

### Genetic admixture and structure

To identify the genetic clusters and infer admixture proportions between individuals, we analyzed the admixture of 67 genomes of Kronotskoe-Uzon charrs in fastSTRUCTURE (Raj et al., 2014) based on 14,768,687 SNPs. The number of genetic clusters (K) that best represent our genome data and proportional ancestry was estimated via ten replicates for each K from 1 to 15 and the better K was determined using script chooseK.py (additional parameter for fastStructure --full). The analysis was performed in two steps: first on the entire dataset, and then on the nosed morph, the Krokur charr and the Kronotskaya River Dolly Varden (24 genomes), which did not split up in the first step.

Per-population observed heterozygosity was estimated from the Stacks ‘populations’ output using variant and invariant positions, providing a measure of individual-level genetic diversity within each population. Genome-wide diversity and divergence landscapes were characterized with pixy (v2.0.0; Korunes & Samuk, 2021) using the populations.all.vcf (all sites) as input, which calculates unbiased estimates of nucleotide diversity (π), absolute divergence (dxy), and relative divergence (*F*_ST_) in non-overlapping 1 Mb windows while accounting for missing data (--stats pi fst dxy, --window_size 1000). Mean and standard deviation of each statistic were summarized across windows for each population (π) and each population pair (dxy, *F*_ST_) using custom awk scripts, to characterize overall patterns of diversity and differentiation among groups. Fish no. 9h (a putative hybrid exhibiting a substantial mixture of two genetic clusters) was excluded from these analyses.

### Historical gene flow

We used Patterson’s D statistics from DSUITE (v0.4; Malinsky et al., 2021) to test for historical gene flow between the Kronotskoe-Uzon charrs. This method is a widely used and robust tool to detect introgression between closely related groups and to distinguish it from incomplete lineage sorting.

Putative hybrid no. 9h s was excluded from the analysis. The consensus tree from IQ-TREE was used with the “d-trios” function to calculate D-statistics for all possible trios. The sample set was used with SNPs from the loci genotyped in at least 80% of the considered samples and with a minor allele count of two, resulting in 2,750,712 SNPs. The significance of introgression events was assessed with the jackknife approach after adjusting for multiple comparisons (Durand et al., 2011). The number of blocks used in the jackknife procedure (--JKnum parameter) was tested in the range from 20 to 60. The results were visualized as a heatmap using the dtools.py script from DSUITE.

### Morphometric comparison

Our previous detailed morphometric analysis allowed to reliably distinguish all Lake Kronotskoe ecomorphs (Esin et al., 2020; 2024). The widehead, nosed and deepwater morphs can easily be distinguished by their head shape and mouth size. In this study, we identified trends in body and head shape shifts using a smaller number of landmarks. The fish were photographed in orthogonal projection with straightened fins, and 14 landmarks were gathered for each photo using TPSDig (v2.16; Rohlf, 2010). The samples of the putatively ancestral ecomorphs and derived populations, along with the outgroup, which were identified based on SNP phylogeny, were analyzed.

The samples were compared in MorphoJ (v1.08; Klingenberg, 2008) via Canonical Variate Analysis (CVA), after generalized Procrustes superimposition. To exclude the impact of fish size on group discrimination, we analyzed the Pearson correlations (r^2^) between the individual CV scores and maximum standard length. To determine the reliability of group discrimination by CVA, we performed a classification analysis (predicted vs. observed classes matrix) for the CVA root scores in StatSoft (v11.0; Hill & Lewicki, 2006). Then, a permutation test against the phylogenetic signal from the SNP-based ML tree was conducted in MorphoJ. The weighted squared-change parsimony tree based on Procrustes coordinates (number of randomization rounds = 10,000) was compared with the phylogenetic ML tree. The differences in head and body shape along the canonical roots (CV) were visualized using a wireframe matrix, which was then post-processed in CorelDRAW v21 to align the analyzed landmarks with real fish outlines.

## Results

### Phylogeny and differentiation of charrs

The best-fit nucleotide substitution model TVMe+R3+ASC supported the monophyly of the Kronotskoe-Uzon charrs, grouping them together in a single cluster sister to the cluster of the populations from other Kamchatkan waterbodies (**Fig. 2a**; log-likelihood of the consensus tree = -5.51 × 10^-6^). The latter diverged into three clades, each corresponding to a different sampled river. The Kamchatka River clade was basal, while the clade of the west-coast population from the Bol’shaya River was the most distant. Within the Kronotskoe-Uzon charr assemblage, the anadromous Dolly Varden from the Shumnaya River was the first to branch. The Kronotskaya River Dolly Varden was found to be in a sister position relative to the nosed morph. Among the Lake Kronotskoe charrs, the ML tree supported the differentiation of lineages corresponding to the predatory, nosed and deepwater charrs with 100% bootstrap support.

**Figure 2.**
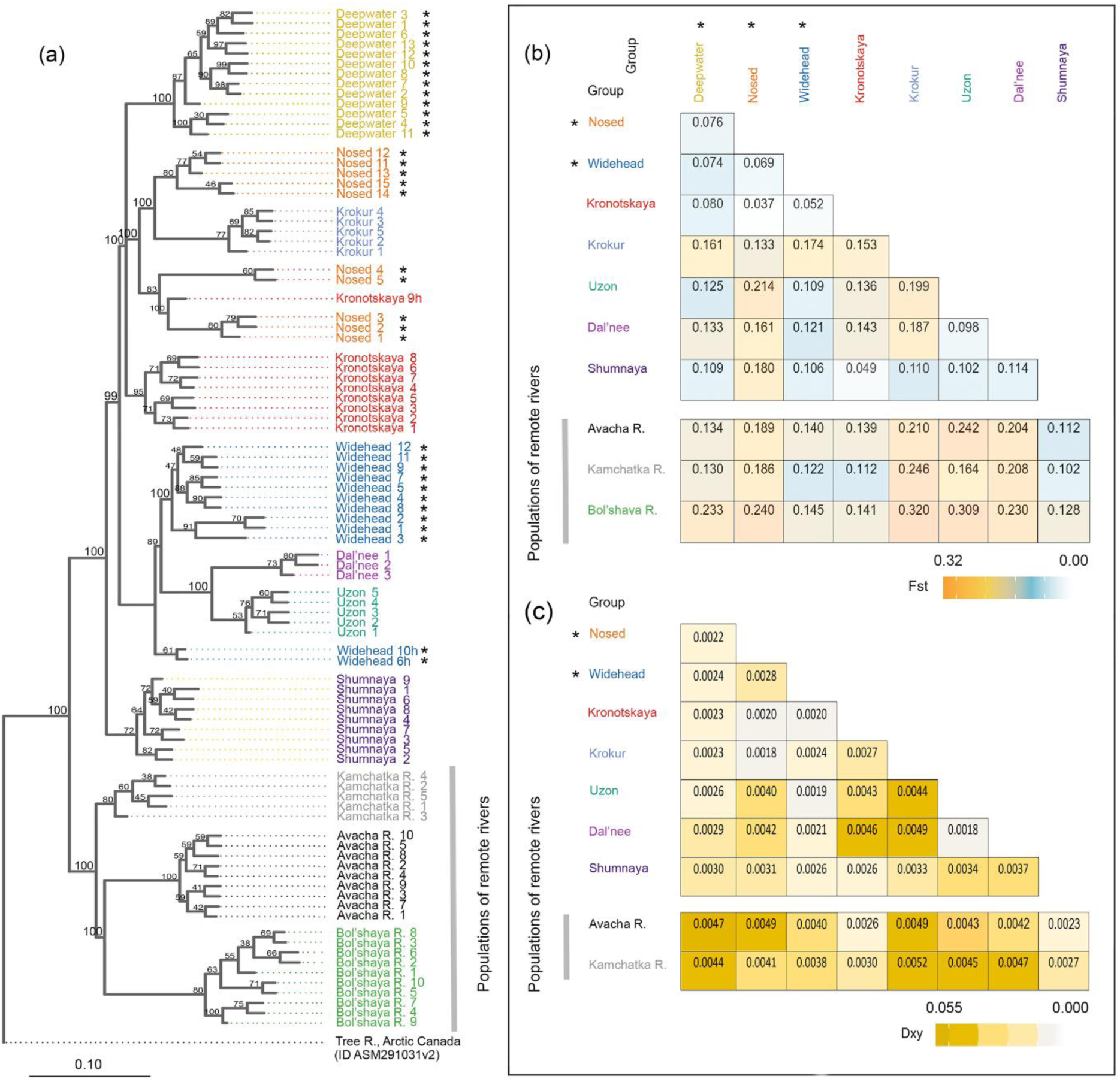
Phylogeny and differentiation of the Kronotskoe-Uzon charrs based on 17,772,739 SNPs. (**a**): Maximum likelihood tree generated from the TVMe+R3+ASC substitution model for individual genomes (numbered as submitted to ENA). The bootstrap support values for each node are shown; the scale is in nucleotide substitutions per site. (**b**): Pairwise *F*_ST_ values and (**c**): Pairwise d_xy_ values among the charr groups. Sample no. 9h was excluded from the analyses (b) and (c). The charr groups are labelled as shown in Figure 1; * indicate Lake Kronotskoe ecomorphs.

Two populations isolated in the Uzon waterbodies (the Uzon and Dal’nee charrs) grouped with the predatory widehead morph. The isolated Krokur charr formed a highly supported monophyletic subclade within the clade of the nosed. Furthermore, one individual from the Kronotskaya River (no. 9h) was found to be grouped with the nosed morph, and two widehead individuals (no. 6h and 10h) fell into a basal short branch within the clade of the widehead morph.

All *F*_ST_ pairwise comparisons were statistically significant, including those of sympatric Lake Kronotskoe morphs, with values ranging from 0.043 (*p* = 0.022) to 0.240 (*p* < 0.001) (**Fig. 2b**). Within the Kronotskoe-Uzon assemblage, the charrs isolated in the Uzon waterbodies and Lake Krokur were the most differentiated. The Lake Dal’nee population was the least different from the Uzon charr, whereas the Lake Krokur population was the least different from the nosed morph. The Kronotskaya River Dolly Varden (including fish no. 9h) showed a relatively weak differentiation, being closer to the Lake Kronotskoe nosed morph (*F*_ST_ = 0.043) than to the Dolly Varden from the Shumnaya River (= 0.084).

All Kronotskoe-Uzon charrs were highly significantly differentiated from the populations of the remote rivers, with the Kamchatka River population from the closest coastal bay being the least divergent.

The d_xy_ estimates within the Kronotskoe-Uzon assemblage showed minimal nucleotide divergence between the Uzon populations and the widehead morph, as well as between the Krokur charr and the nosed morph. On average, these groups diverged from each other by 0.18‒0.21% substitutions, whereas sympatric lacustrine lineages diverged by 0.22‒0.28% substitutions (**Fig. 2c**). The greatest distances (≥0.45%) were observed among the Dal’nee charr ‒ Krokur charr ‒ Kronotskaya River Dolly Varden, as well as between the Kronotskoe-Uzon charrs and the Dolly Varden from the remote rivers.

### Genetic structure and hybridization of charrs

The optimal number of genetic clusters within the Kronotskoe-Uzon assemblage was found to be four: log-likelihood = -1.165 × 10^-8^ at K = 4 and ≤ -1.172 × 10^-8^ at another K. The admixture analysis differentiated the Shumnaya River Dolly Varden (i), the lineage of Lake Kronotskoe deepwater morph (ii), the lineage of predatory widehead morph and the Uzon charr (iii) and the lineage of nosed morph combined with the Krokur charr and Kronotskaya River Dolly Varden (iv) (**Fig. 3a**). Two individuals of the widehead morph (№ 6h and 10h) exhibited a mixture of the second and third clusters.

**Figure 3.**
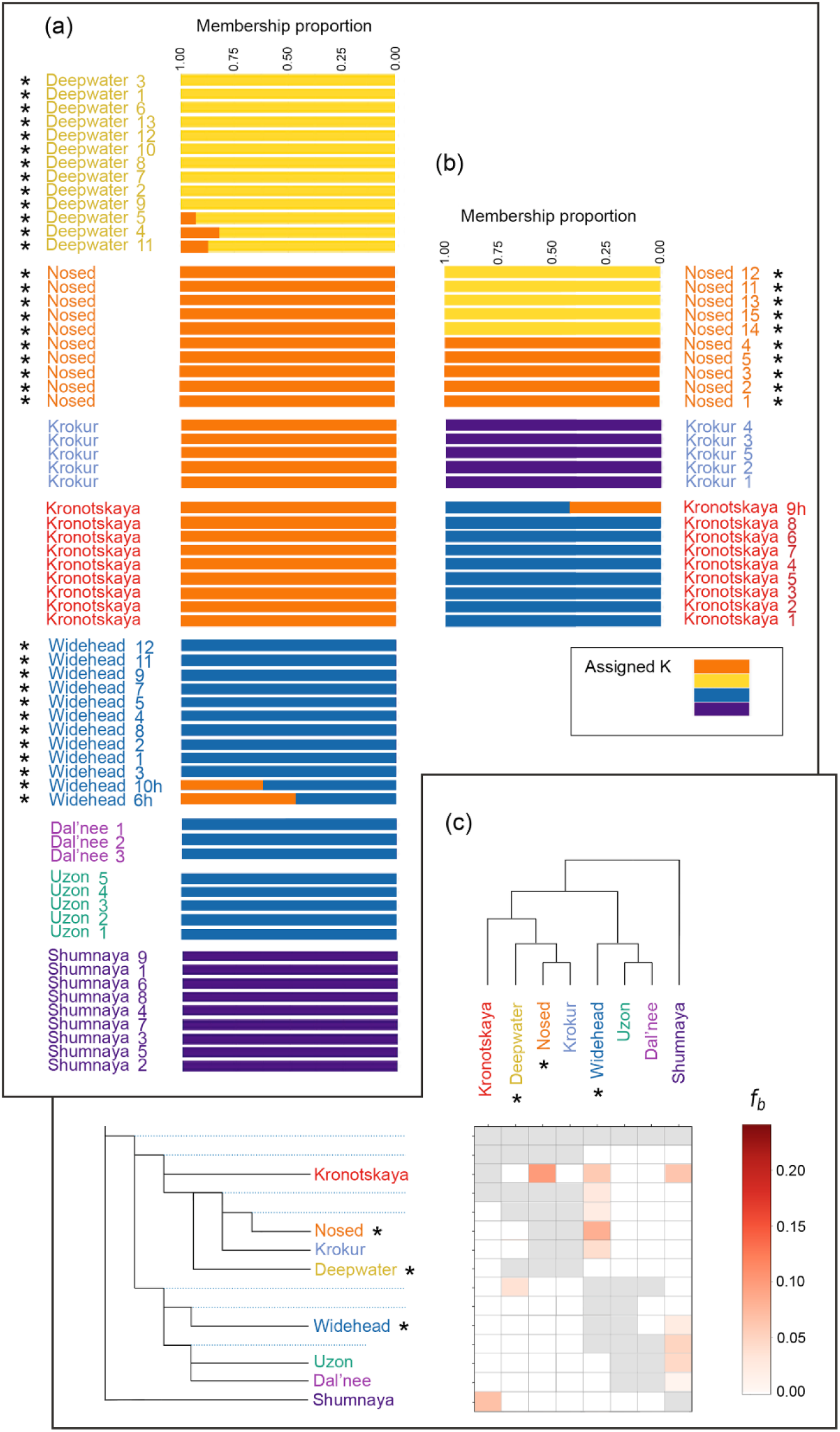
Genetic structure and hybridization of the Kronotskoe-Uzon charrs. (**a** and **b**): The current genetic admixture presented as bar plots of the genetic cluster proportions (coloured) inferred by two consecutive steps of the fastSTRUCTURE analysis based on 14,768,687 SNPs. The statistically optimal number of genetic clusters for 67 (a) /24 (b) genomes based on log-likelihood estimates is 4/4. (**c**): A matrix of inferred historical gene flow as estimated from gene trees for Kronotskoe-Uzon populations employing 2,750,712 SNPs. The matrix shows the inferred f-branch metric, reflecting excess allele sharing between the branch of the ‘laddered’ full-phylogenetic tree on the Y-axis (relative to its sister branch), and the branches defined on the X-axis. The dotted lines of the full-phylogenetic tree represent inferred ancestral genotypes. Grey cells indicate comparisons that cannot be made owing to the tree topology, white cells represent no evidence of gene flow. Sample no. 9h was excluded from the analysis (c). The charr groups are labelled as shown in Figure 1;* indicate Lake Kronotskoe ecomorphs.

In the second step of the analysis, the Lake Krokur charr (i), the Kronotskaya River Dolly Varden (ii) and two clusters of the nosed morph (iii and iv) were differentiated (**Fig. 3b**; log-likelihood = -1.122 10^-8^ at K = 4 and ≤ -1.130 10^-8^ at another K). FastSTRUCTURE also indicated the admixture of one of the nosed charr clusters to the fish no. 9h, which was caught in the Kronotskaya River downstream rapids.

The genome nucleotide diversity (π) and heterozygosity (Ho) of the landlocked Kronotskoe-Uzon charrs was found to be lower than those of the anadromous Dolly Varden populations (**Table 1**). The Lake Kronotskoe ecomorphs exhibited higher diversity and heterozygosity than the derived Uzon and Krokur populations. The populations from the maar lakes exhibited the lowest genetic variability. The π and Ho values based on SNPs were fully consistent with the previously estimated genetic diversity and heterozygosity based on STR and mtDNA markers (Esin et al., 2025).

**Table 1.**
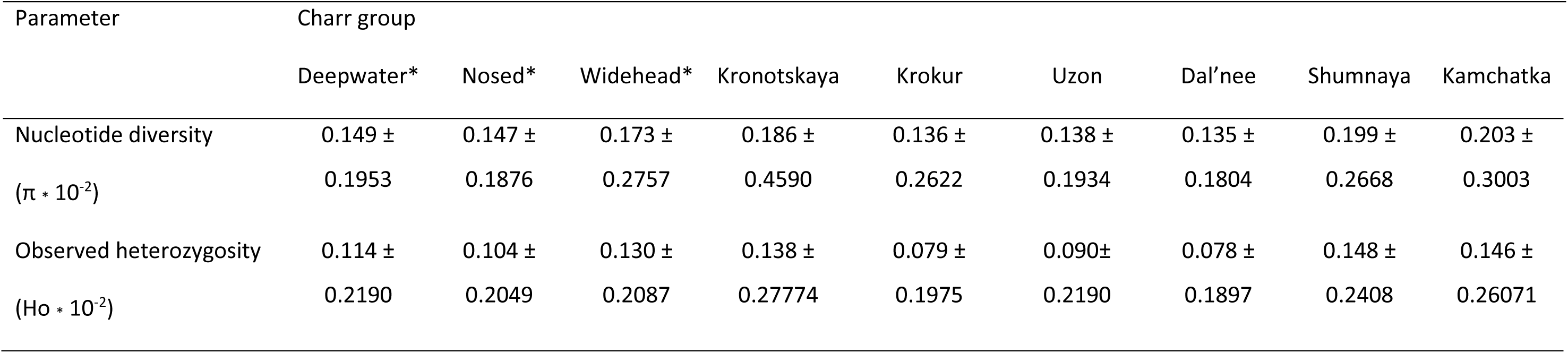
Genetic diversity and heterozygosity (mean ± Std. Dev) of the Kronotskoe-Uzon charrs and the Kamchatka River Dolly Varden (* indicate Lake Kronotskoe ecomorphs)

| Parameter | Charr group |  |  |  |  |  |  |  |  |
| --- | --- | --- | --- | --- | --- | --- | --- | --- | --- |
|  | Deepwater* | Nosed* | Widehead* | Kronotskaya | Krokur | Uzon | Dal'nee | Shumnaya | Kamchatka |
| Nucleotide diversity | 0.149 ± | 0.147 ± | 0.173 ± | 0.186 ± | 0.136 ± | 0.138 ± | 0.135 ± | 0.199 ± | 0.203 ± |
| ($\pi \cdot 10^{-2}$) | 0.1953 | 0.1876 | 0.2757 | 0.4590 | 0.2622 | 0.1934 | 0.1804 | 0.2668 | 0.3003 |
| Observed heterozygosity | 0.114 ± | 0.104 ± | 0.130 ± | 0.138 ± | 0.079 ± | 0.090± | 0.078 ± | 0.148 ± | 0.146 ± |
| ($H_o \cdot 10^{-2}$) | 0.2190 | 0.2049 | 0.2087 | 0.27774 | 0.1975 | 0.2190 | 0.1897 | 0.2408 | 0.26071 |

After excluding fish no. 9h with admixed ancestry, 78 significant tests of trio combinations were compiled using D statistics across the eight analyzed groups of the Kronotskoe-Uzon assemblage. Among 13 gene flow events detected, admixture levels of over 5% were estimated for seven of them using the f-branch metric (**Fig. 3c**). Two of these events involved gene flow from the predatory and nosed morphs of Lake Kronotskoe to the Dolly Varden of the Kronotskaya River. The inferred hybridization matrix also demonstrated gene flow between the lineages of predatory and nosed (including the Krokur derivative) charrs, which is in accordance with the data on contemporary hybridization revealed by fastSTRUCTURE (putatively hybrid fish no. 6h and 10h). Weak historic gene flow was detected between the Dolly Varden from the Kronotskaya and Shumnaya rivers. Finally, traces of hybridization were evaluated between the Shumnaya River Dolly Varden and the Uzon charr.

### Morphometric specifics of charrs

CVA demonstrated that the first three canonical roots accounted for 87.6% of the variation in morphometric traits. Herewith, CV1 (43.6% of variance) mainly described allometric differences, with r^2^ = 0.62 for the CV1 score‒standard length ratio. In contrast, the CV2 and CV3 scores were weakly correlated with fish length (= 0.09 and 0.01, respectively), allowing their use to determine patterns of group differentiation in body and hade shape (**Fig. 4a**). We found that CV2 primarily separated the populations from the maars Krokur and Dal’nee, while CV3 significantly contributed to separating the widehead and nosed morphs from Lake Kronotskoe. The CV2‒CV3 morphospace allowed us to distinguish the maar populations from lakes Krokur and Dal’nee, as well as the two endemic ecomorphs from Lake Kronotskoe; the convex hull areas of the Uzon charr and the Kronotskaya River Dolly Varden were joint and adjacent to the area of Kamchatka River Dolly Varden.

**Figure 4.**
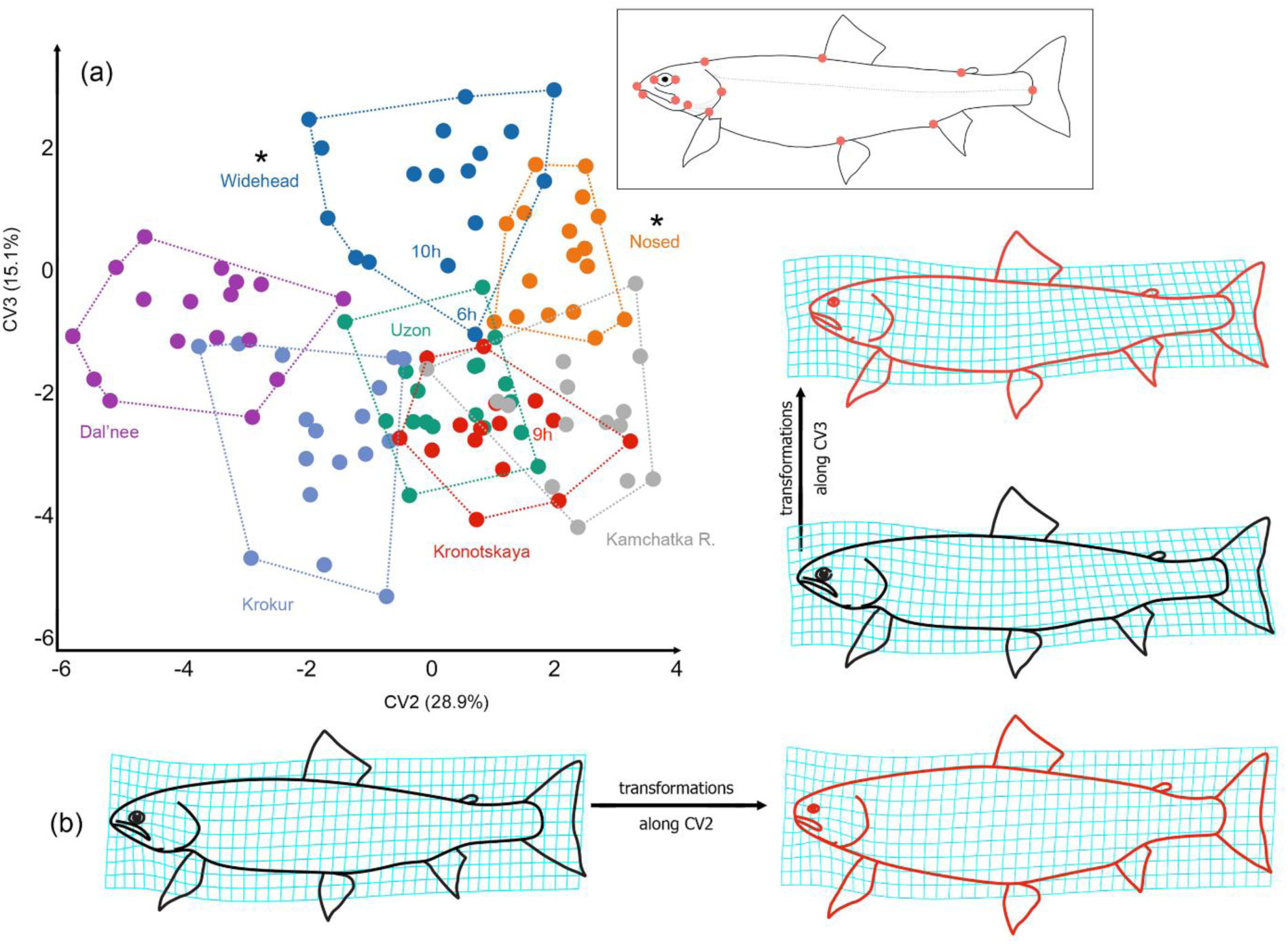
Results of the morphometric comparison of the Kronotskoe-Uzon charrs derived from canonical variate analysis, with the anadromous Dolly Varden from the Kamchatka River as an outgroup. (**a**): Body shape comparison in CV2‒CV3 space (% of total variance in brackets), convex hull areas for each group are shown; inset to the right represents landmarks’ position used in the morphometric analysis. Additionally, the three individuals with admixed ancestry are indicated with numbers. (**b**): Deformation of the grids representing the change in body shape along the second and third canonical roots. The charr groups are labelled as shown in Figure 1;* indicate Lake Kronotskoe ecomorphs.

Grid deformations indicated head shortening, eye diameter and mouth size decrease, and body depth increase along CV2, as well as head height increase and snout shortening along CV3 (**Fig. 4b**).

To validate the CVA outcomes, we performed a classification analysis of the CV2‒CV6 scores. The algorithm correctly classified the Krokur charr (no misclassifications), the Dal’nee charr (one misclassification with the widehead morph), the predatory widehead morph (one misclassification with the nosed morph and one with the Dal’nee charr) and the nosed morph (one misclassification with the Uzon charr and one with the widehead morph). One out of three fish (no. 6h), exhibited a specific phylogenetic position and genetic admixture, was misclassified within the widehead morph sample (**Fig. 4a**). The classification failed for the Uzon charr, the Kronotskaya River Dolly Varden and the Kamchatka River Dolly Varden because the algorithm grouped them all together. The Dolly Varden no. 9h was found within the convex hull of the Kronotskaya River sample.

A weighted squared-change permutation test based on Procrustes coordinates revealed no correlation between the pattern of group morphological differentiation and their phylogenetic relationships (test *p* = 0.202). Therefore, no phylogenetic signal was evident in the shifts in head and body shape.

## Discussion

The contribution of endemic fish ecomorphs (species) originating from local adaptive radiations to shaping the regional and global biodiversity remains a subject of ongoing discussion. There are only a few examples of the secondary expansion of ecomorphs beyond their cradle ecosystems (Daufresne et al., 2004; Kullander et al., 2011; Levin et al., 2019; Yousefi et al., 2020; Sideleva et al., 2025). This study reveals the different ways in which postglacial adaptive morphs of salmonids disperse from the landlocked ecosystem into neighbouring basins. Derived populations appear to lose the specific morphological characteristics of the ancestral lineages, instead acquiring generalized morphotypes appropriate to lacustrine or riverine habitats. In modern conditions, the derivatives do not expand further, and this pattern of endemic spreading only enriches biodiversity on a local scale.

We found that the descendants of *S. malma* formed a local gene pool in the Kronotskoe-Uzon catchment of the late Pleistocene age due to evolution under insufficient gene flow from surrounding Kamchatkan populations. Analysis of SNP data confirms the radiation of endemic charrs from Lake Kronotskoe into three phylogenetic lineages. They differ by an average of 0.20‒0.24% substitutions and may occasionally hybridize, as determined by our analyses of genetic structure and admixture. However, the lineages are genetically differentiated. Our genomic data suggest that the two lineages represented by predatory and littoral benthivorous charrs invaded the newly formed peripheral waterbodies of the Holocene age and continued to migrate downstream from Lake Kronotskoe, hybridizing with the anadromous Dolly Varden in the outflowing river. These dispersal processes gave rise to three new isolates and one hybrid population.

We found that the ancestor of the predatory widehead morph from Lake Kronotskoe populated the Shumnaya River in the Uzon caldera via a temporary waterway. On the phylogenetic tree, the charr from the Uzon caldera is grouped with the endemic predator rather than with the Dolly Varden from the lower course of the Shumnaya River or with other populations. The admixture test also indicates the shared ancestry of the Uzon charr and the widehead morph. The predatory charr could have colonized the Uzon caldera at the Pleistocene-Holocene boundary from the Zelenaya River headwaters (**Fig. 5a** and **b**). Glaciers were well-developed in this region (Braitseva et al., 1995; Golub, 2006), resulting in the release of substantial volumes of meltwater following climate warming. Numerous traces of the ancient dried riverbeds can be found on the plateau between Lake Kronotskoe and the Uzon caldera. This riverbed pathway was disrupted by a catastrophic landslide that occurred 8,500 years ago (Ponomareva et al., 2006). The Lake Kronotskoe predatory charr still reproduces in the Zelenaya River headwaters (Markevich et al., 2021), whereas the Uzon charr migrates to spawn in the Zapadny stream (**Fig. 5b**). The converging tributaries of Lake Central’noe and the ancient maar Dal’nee in the Uzon caldera allowed the Uzon charr to populate the maar (**Fig. 5b**). Our analyses confirm the close kinship of the Uzon and Dal’nee charrs. High *F*_ST_ and low d_xy_ between the Uzon charrs and the widehead morph, as well as reduced nucleotide diversity and heterozygosity in the derived populations compared to the widehead morph suggest that only a small number of charrs initially populated the caldera waters.

**Figure 5.**
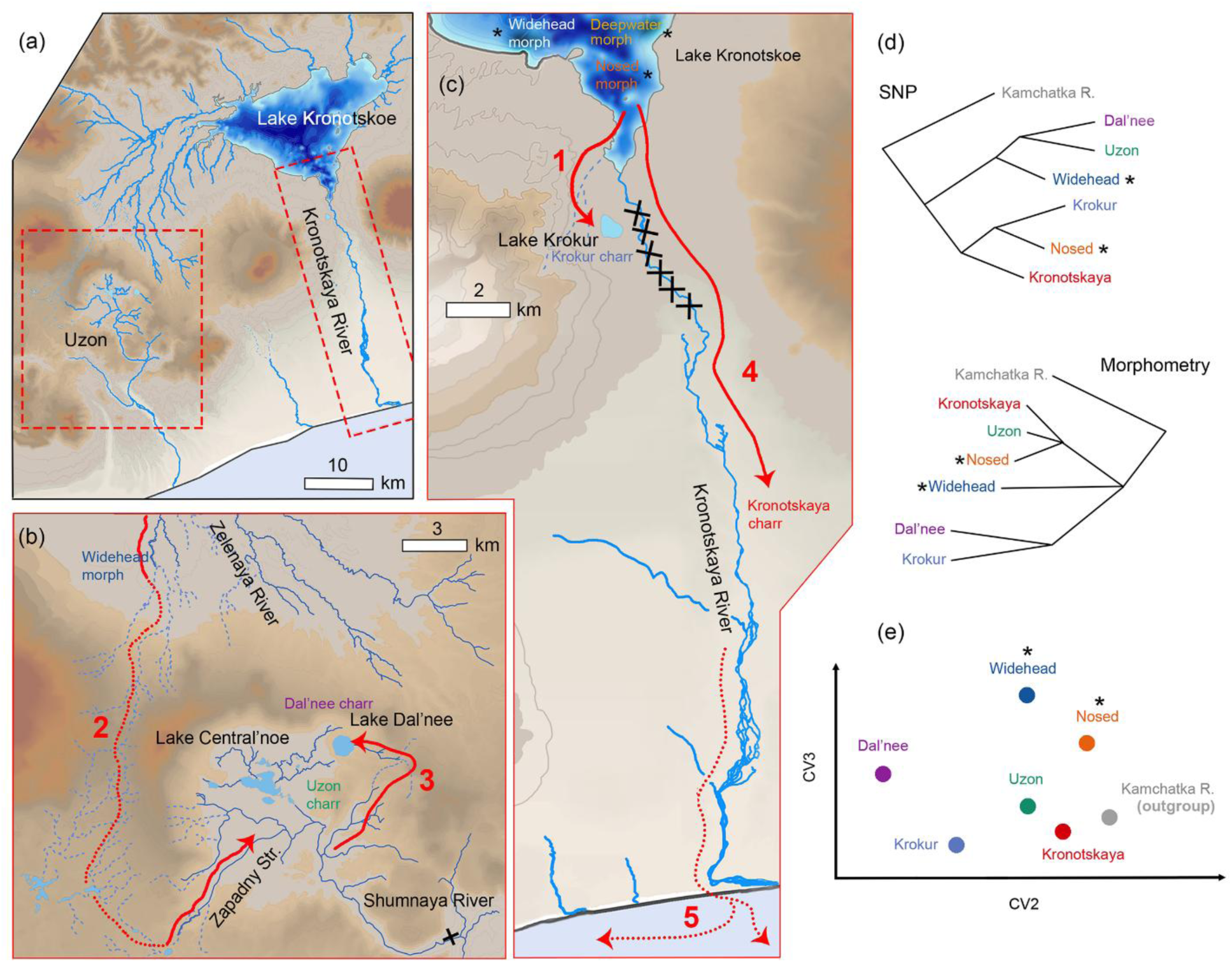
Scheme showing the expansion of the Lake Kronotskoe charrs. (**a)**: General view of the catchment area. (**b** and **c**): Migration routes of charrs into the Uzon caldera and downstream the basin, respectively. The arrows on the maps denote the pathway of the endemic charrs’ settlement into Lake Krokur (1), the Uzon caldera catchment (2), Lake Dal’nee (3), the lower course of the Kronotskaya River (4) and the secondary dispersal of hybrids through the marine environment (5); symbols: x ‒ waterfalls and rapids, ‒ temporary watercourses. (**d**): Сontraposition of the phylogeny of Kronotskoe-Uzon charrs derived from a maximum likelihood approach based on SNPs and a Procrustes coordinate weighted squared-change parsimony approach. (**e**): Representation of the group centroids’ position in the canonical variate morphospace derived from CVA. The charr groups are labelled as shown in Figure 1; * indicate Lake Kronotskoe ecomorphs.

Expanding the range through temporary water interconnections is a common phenomenon among fish. For instance, the endemic *Labeobarbus* spp. were expected to resettle across river headwaters in the Abyssinian highlands (Levin et al., 2019; 2020), while *Thymallus* spp. resettled across rivers in southern Siberia (Weiss et al., 2007). However, despite the well-developed connections between the sources of the Penjina and Anadyr rivers (Koval et al., 2018), the endemic whitefish species *Coregonus subautumnalis* from the Penjina River and *C. anaulorum* from the Anadyr River were unable to invade each other’s habitats. This highlights the importance of migratory activity in the potential for expansion in fish. No descendants of the deepwater charrs from Lake Kronotskoe were found in the surrounding basins, which supports the hypothesis that this lineage has limited migratory ability (Markevich et al., 2021). Our phylogenetic analyses and genetic divergence estimates also do not suggest a close relationship between the Lake Kronotskoe endemics and the Kamchatka River Dolly Varden. Probably, predatory and nosed charrs were unable to overcome the steep watersheds leading to the Kamchatka River basin. Even if resettlement had once taken place, the small number of migrants did not produce any genetic signal.

At the end of the Pleistocene, the Uzon caldera was locked and filled with a periglacial lake (Egorova, 1993; Bazhenova et al., 1998). In the early Holocene, a catastrophic landslide occurred on the south-eastern side of the caldera (Leonov et al., 1991), resulting in the range break, the draining of the lake, and the formation of a canyon with a 50 m waterfall on a lava ledge. Judging by the signal of historical hybridization with the Uzon charr, we hypothesize that some Dolly Varden accessed the caldera reservoir prior to the formation of the waterfall. Alongside this, the phylogenetic tree topology, high genetic divergence and undetected admixture all suggest limited present-day contact between the Uzon charr and the Shumnaya River Dolly Varden. It appears that charrs attempting to run downstream die after falling over the waterfall. The height of the 50-m gorge is sufficient to cause critical injury to fish (Pavlov & Skorobogatov, 2014).

The drainless maar Krokur originated around 6,000 years ago (Ponomareva et al., 2017) seems to be stocking from a now-disappeared tributary of Lake Kronotskoe, which was located where the maar erupted. Following the cold phreatic explosion, some fish may have survived in the tributary section on the crater slope and colonized the reservoir (**Fig. 5a** and **c**). There are no watercourses that could have provided an alternative pathway for the fish invasion. A similar process is thought to account for the endemic *Amphilophus* cichlids having settled in the Nicaraguan drainless crater Xiloa (Kusche et al., 2014), and *Procatopus* poecilids and *Barbus* cyprinids having invaded the crater lake Dissoni near the Uve River in Cameroon (Schliewen et al., 1994; Schliewen, 2005). The topology of the phylogenetic tree, d_xy_ estimates and the shared ancestry revealed by admixture tests strongly support that the Krokur charr originated from a littoral benthivorous (nosed) ancestor. Small brooks in the basin, such as a former tributary on the maar site, are commonly used for reproduction exactly by the nosed morph (Markevich et al., 2021). As with the Uzon charr, the Lake Krokur derivative population exhibits high *F*_ST_ and reduced genetic diversity and heterozygosity compared with the nosed morph, indicating its origin from a small initial group.

The third dispersal scenario appears to have occurred once the rapids on the Kronotskaya River became passable downstream. Endemic ecomorphs migrate from Lake Kronotskoe to the reproduction zone of the Kronotskaya River Dolly Varden (= Kronotskaya charr) (**Fig. 5c**). Our analysis revealed that the riverine population is genetically distinct from the anadromous *S. malma* inhabiting the surrounding rivers, being a part of the endemic Kronotskoe-Uzon assemblage. D-statistics indicate that gene flow has occurred from the endemic Lake Kronotskoe predatory and nosed morphs to the riverine population.

Using fastSTRUCTURE, we identified an individual from the Kronotskaya River lower course with the substantial admixture from the nosed morph, suggesting limited ongoing hybridization downstream of the rapids. The Kronotskaya charr, which migrates to the ocean in summer (Esin et al., 2020; 2025), differs in the highest genetic diversity and heterozygosity among the assemblage members. The noticeable genetic difference observed between the Kronotskaya charr and the population of the nearby Shumnaya River, coupled with the phylogenetic separation of the Kronotskaya charr from remote anadromous populations, support the previously mentioned reproductive philopatry among Kamchatkan charrs (Markevich et al., 2021). It seems that, regardless of their origin, anadromous populations only return to spawn in their native rivers in modern conditions.

Our morphometric analysis revealed that the main vectors of charr morphological divergence emerged independently of the morphs’ phylogenetic history. The groups analyzed did not branch in the same way on the morphometric and phylogenetic trees (**Fig. 5d**). The group centroids positioned in the canonical variate morphospace were divided into three sectors (**Fig. 5e**). The left sector comprised the flooded crater populations, the upper sector consisted of the Lake Kronotskoe adaptive morphs and their putative hybrids, and the bottom-right sector comprised groups with the generalized Dolly Varden morphotype, including the Kamchatka River Dolly Varden, which had no contact with the Kronotskoe-Uzon assemblage according to SNP data.

After settling in flooded maar craters with similar environmental conditions, the descendants of two phylogenetic lineages demonstrated similar morphological changes. Pelagic fish from the maars Krokur and Dal’nee have an elongated head, sharp snout and low body. These features distinguish them from the robust-bodied demersal representatives of their evolutionary lineages in Lake Kronotskoe. Similar morphological transformations associated with resettlement between demersal pelagic habitats are evident in the endemic Transcaucasian scrapers of the genus *Capoeta* (Levin et al., 2012; Pepoyan et al., 2014) and Tibetan snowtrouts *Schizothorax* spp. (Regmi et al., 2021). Regarding the Uzon charr, it may be hypothesized that this ecomorph has experienced the effect of ecological release in the functional morphology (Herrmann et al., 2021) due to a shift in lifestyle from lacustrine-addfluvial to primarily riverine, as well as the loss of predatory ancestor trophic specialization (Esin et al., 2015). It has become more similar in appearance to the generalized *S. malma*.

It can be assumed that the settlers initially possessed specific morphological characteristics, since the colonization of peripheral waterbodies occurred after the diversification of the main adaptive lineages of Lake Kronotskoe charrs. The decrease in initial variability, which was potentially due to the founder effect, did not prevent the formation of a new morphotype suitable for the new habitat in all three derivatives. This is reminiscent of how specialized trunk-ground anole lizards change their morphology to an arboreal lifestyle after multiple inter-island resettlements (Losos & Queiroz, 1997). The question of whether such morphological transformations result from rapid selection or whether a new suitable variant emerged from reserved plasticity of the ancestral populations remains to be resolved.

Our findings emphasize the importance of the continuous pressure of local environmental factors in the forming and maintaining specific adaptations within a particular food niche in a given radiation. However, since these factors may vanish in novel ecosystems, morphological adaptations could rapidly collapse and switch to more appropriate features. This also shows the futility of morphological analysis solely in reconstructing the phylogeny of ecologically plastic fish species that produce contrasting phenotypes in different habitats. A defining feature of adaptive radiation is the accelerated diversification of phenotypes relative to the gradual genetic assimilation (Schneider & Meyer, 2017; McGee et al., 2020).

Despite hybridizing with endemic ecomorphs, the Kronotskaya charr exhibits no striking morphological features and visually resembles Dolly Varden from the Kamchatka River. Nevertheless, we believe that the population of the Kronotskaya River differs fundamentally from the maar isolates in terms of its evolutionary prospects. The isolates can only produce new ecomorphs after the water network reorganization. Their genetic diversity and heterozygosity have been reduced, whereas the genetic diversity and heterozygosity of the Kronotskaya charr are the highest in the assemblage. A generalized group with a genome saturated with “introduced alleles” is likely to acquire increased ecological plasticity and gained advantages in dispersal during environmental changes. An increase in genome heterogeneity was found to contribute to the ecological diversification and speciation of cichlids (Meier et al., 2017; Irisarri et al., 2018; Svardal et al., 2020). Gene transfer from specialized resident populations to anadromous *Gasterosteus aculeatus* enabled the latter to rapidly form new adaptive morphs when invading new freshwater habitats (Schluter & Conte, 2009). The same evolutionary mechanism was also suggested to explain rapid diversification of European *Coregonus* spp. (Hirsch et al., 2013), *Labeobarbus* spp. of the Abyssinian highland waterbodies (Levin et al., 2024), and American *Crenicichla* cichlids (Burress et al., 2023). In these cases, the re-assembly of genetic variation into new combinations may have facilitated the diversification and speciation over short time periods (Marques et al., 2019; McGee et al., 2020). *Salvelinus malma* is a remarkable example of these processes in Kamchatka (Esin, 2025).

To conclude, our findings suggest that locally radiated postglacial ecomorphs of salmonids can migrate from the cradle ecosystem and successfully populate surrounding waterbodies, despite their strict adaptations to particular ecological niches. The variety of identified expansion patterns demonstrates the universality of the process and the potential for its replication in other areas of adaptive radiation. Ecomorph descendants evolve new morphotypes in response to changes in environmental pressures over a period of time insufficient for their genetic ancestry to be blurred. The downstream migration of adaptive morphs followed by their hybridization with the population of ancestral type is likely to be pivotal in enabling local endemics to contribute to biodiversity enrichment at the regional level.

## Acknowledgements

We would like to express our deep gratitude to Dr Anastasia Teterina (University of Oregon), who processed the bioinformatic data. We also thank Dr Boris Levin (IEE RAS) for useful criticism of the results. We are grateful to Dr Daria Panicheva (Bering KSU) for her invaluable help in organizing the work, which was essential to the project’s success.

## Data Availability Statement

Raw Illumina reads have been submitted to the European Genome-Phenome Archive (ENA) and are available under BioProject study number ERP185516 (samples ERS28138360 … ERS28384601). https://www.ebi.ac.uk/ena/browser/view/PRJEB104211

